# Variability in Locomotor Dynamics Reveals the Critical Role of Feedback in Task Control

**DOI:** 10.1101/764621

**Authors:** Ismail Uyanik, Shahin Sefati, Sarah A. Stamper, Kyoung-A Cho, M. Mert Ankarali, Eric S. Fortune, Noah J. Cowan

## Abstract

Animals vary considerably in size, shape, and physiological features across individuals, but yet achieve behavioral performances that are virtually indistinguishable between conspecifics. We examined how animals compensate for morphophysiological variation by measuring the system dynamics of individual knifefish (*Eigenmannia virescens*) in a refuge tracking task. Kinematic measurements of *Eigenmannia* were used to generate individualized estimates of each fish’s locomotor plant and controller revealing substantial variability between fish. To test the impact of this variability on behavioral performance, these models were used to perform simulated ‘brain transplants’—computationally swapping controllers and plants between individuals. We found that simulated closed-loop performance was robust to mismatch between plant and controller. This suggests that animals rely on feedback rather than precisely tuned neural controllers to compensate for morphophysiological variability.

## Introduction

Animals routinely exhibit dramatic variations in morphophysiology between individuals but can nevertheless achieve similar performance in sensorimotor tasks (***Sponberg et al., 2015***; ***Bullimore and Burn, 2006***). Further, individual animals can experience rapid changes in their own morpho-physiological features, such as extreme weight changes that occur during and between bouts of feeding. For example, mosquitos can consume more than their body weight (***Van Handel, 1965***) and hummingbirds can consume up to 20 % of their body weight (***Hou et al., 2015***) in a single feeding. How neural control systems accommodate these changes is not known.

The behavioral performance of an individual animal is determined via an interplay between its ‘controller’ and ‘plant’ (***Kiemel et al., 2011***; ***Van Der Kooij and Peterka, 2011***; ***Cowan et al., 2014***; ***Dickinson et al., 2000***; ***Hedrick et al., 2009***). The plant typically includes musculoskeletal components that interact with the environment to generate movement (***Hedrick and Robinson, 2010***; ***Sefati et al., 2013***; ***Maladen et al., 2009***). The controller typically includes sensory systems and neural circuits used to process information to generate motor commands (***Cowan and Fortune, 2007***; ***Kiemel et al., 2011***; ***Lockhart and Ting, 2007***; ***Roth et al., 2014***). From the perspective of control theory, one might expect the dynamics of the controller to be precisely tuned to the dynamics of the plant, resulting in an optimal control law that reduces variability in task performance (***Todorov, 2004***; ***Franklin and Wolpert, 2011***; ***Bays and Wolpert, 2007***). Were this the case, variations across individuals in morphophysiological features of their plants should manifest in commensurate differences in each animal’s controller. Alternatively, the central nervous system (CNS) may be implementing robust feedback control that attenuates morphophysiological variability at the behavioral level without the need for precise tuning.

Investigating this phenomena requires separate estimates for plants and controllers. However, the classical input–output system identification of behavioral tasks—using only the sensory input and the behavioral output—is limited to generating closed-loop control models of behavioral responses. Data-driven system identification of the underlying neural controllers or locomotor plants requires additional observations such as a measurement of the control output. Electromyograms (EMGs) are the most commonly used proxy for the output of the neural controller. EMGs allow separate data-driven estimates of the controller and plant (***Kiemel et al., 2011***; ***Van Der Kooij and Peterka, 2011***) but require understanding the coordination strategy across multiple groups of muscles that interact in fundamentally nonlinear ways (***Ting and Macpherson, 2005***).

We studied refuge tracking in a species of weakly electric fish *Eigenmannia virescens* (***Figure 1***), a system that permits identification of input–output dynamics as well as the locomotor plant via behavioral observations alone. Like an “aquatic hummingbird”, *Eigenmannia* precisely hovers in place, making rapid and nuanced adjustments to its position in response to the movement of the refuge in which it is hiding (***Rose and Canfield, 1993***; ***Roth et al., 2011***; ***Uyanik et al., 2019b***) (***Figure 1***–***video 1***). During refuge tracking, individual *Eigenmannia* generate fore-aft thrust forces using undulatory motions of their ventral ribbon fin. Undulations are initiated at the rostral and caudal ends of the fin resulting in counter propagating waves that travel towards each other (***Sefati et al., 2013***; ***Ruiz-Torres et al., 2013***). The two traveling waves meet at a position along the ribbon fin known as the nodal point (***Figure 2***A, ***Figure 2–Figure Supplement 1, Figure 2–video 1***). In a task in which the fish maintains position in a stationary refuge, it has been shown that they shift the rostrocaudal position of the nodal point as a function of steady-state swimming speed (***Sefati et al., 2013***). Indeed, transient movements of the nodal point during dynamic refuge-tracking behavior can be used as a linear proxy of the output of the neural controller, *u*(*t*). This feature permits separate identification of the controller and plant for a freely moving animal via behavioral observations alone. Using this behavioral model system, we examined the variability of plants and controllers across individuals.

**Figure 1.**
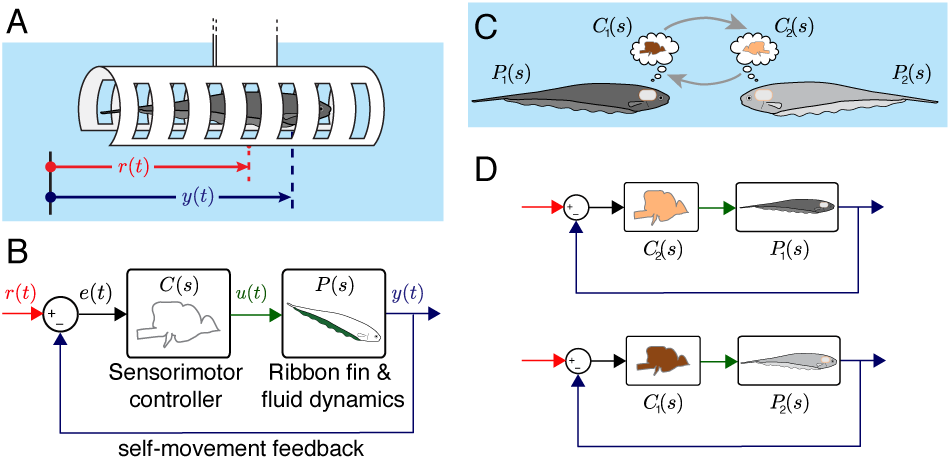
Experimental and computational approach. (**A**) *Eigenmannia virescens* maintains its position, *y*(*t*), in a moving refuge. *r*(*t*) is the position of the moving refuge. (**B**) A feedback control system model for refuge tracking. The feedback controller *C*(*s*) maps the error signal, *e*(*t*), to a control signal *u*(*t*). The plant *P* (*s*), in turn, generates fore-aft movements *y*(*t*). We used measurements of the nodal point as *u*(*t*) to estimate *P* (*s*) in each fish, which was subsequently used to back-calculate *C*(*s*). (**C**) We computationally swapped the individually tailored controllers between fish. (**D**) Feedback control topology illustrating computational brain swaps. The controller from the light gray fish, *C*_2_(*s*), is used to control the plant of the dark gray fish, *P*_1_(*s*), and vice versa. **Figure 1–video 1.** A real-time video recorded at 30 fps from below of an Eigenmannia virescens tracking a moving refuge at 0.55 Hz. The movement of the refuge and fish are plotted below the video.

## Results

We measured the movement of the nodal point of the ribbon fin of *Eigenmannia* in a refuge tracking task. We fit the parameters of a physics-based model (***Sefati et al., 2013***) of ribbon-fin locomotor dynamics to these swimming and nodal point data. Next, we inferred the complementary neural controllers for each individual by using a feedback control model of the input–output behavioral performance (***Cowan and Fortune, 2007***). Finally, we performed computational analysis to assess the role of plant and controller dynamics and feedback in achieving robust behavioral performances.

### Estimating a Data-Driven Plant Model

We measured the positions of the nodal point *u*(*t*), refuge *r*(*t*), and fish *y*(*t*) during refuge tracking. We found that there is an approximately linear relationship between the movement of the nodal point and the velocity of the fish (***Figure 2, Figure 2–Figure Supplement 2***). Hence, we used the nodal point movements as a linear correlate of the output of the neural controller, *u*(*t*), that drives the locomotor plant.

**Figure 2.**
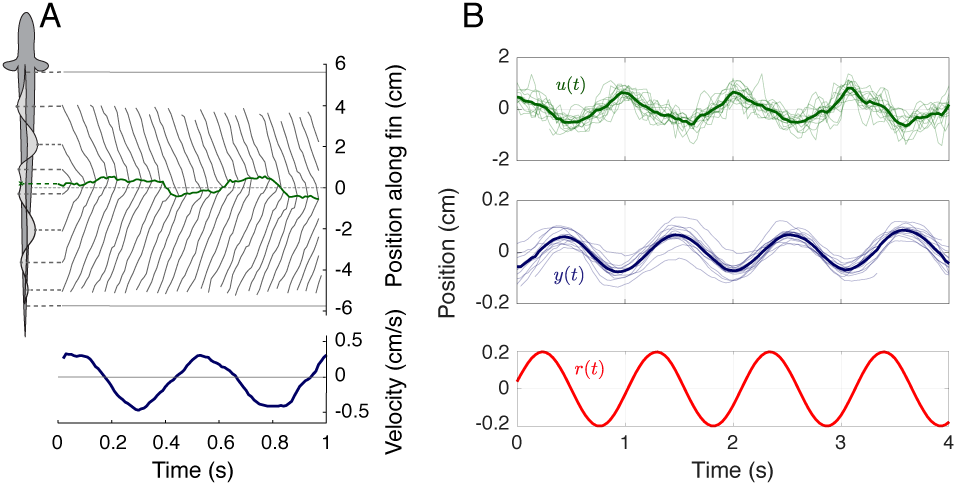
Movement of the nodal point during refuge tracking. (**A**) *Eigenmannia virescens* partitions its ribbon fin into two counter-propagating waves that generate opposing forces (***Sefati et al., 2013***; ***Ruiz-Torres et al., 2013***). The rostral-to-caudal and caudal-to-rostral waves collide at the nodal point (green line). Movements of the nodal point serve as a proxy for the control signal, *u*(*t*). Gray lines indicate the positions of the peaks and troughs of the traveling waves over time. In this trial, the refuge was following a sinusoidal trajectory at 2.05 Hz. (**B**) Movement of the nodal point *u*(*t*), position of the fish *y*(*t*), and position of the refuge *r*(*t*) of 13 trials at 0.95 Hz. Semi-transparent lines represent data from individual trials, green and blue lines are the means. **Figure 2–video 1.** A high-speed video recorded at 100 fps from below an Eigenmannia virescens while tracking a moving refuge at 2.05 Hz. The video is stabilized with respect to fish position to illustrate the movements of the head and tail waves that form the nodal point. The inset shows the rectangular area around the ribbon fin of the fish. The video is played in real-time (10 repetitions), at 0.3× speed (3 repetitions) and at 0.1× speed (1 repetition).

We estimated the frequency response functions (FRFs) of the locomotor dynamics of each fish using the position of the fish *y*(*t*) and its nodal point *u*(*t*). Nyquist plots of the estimated FRFs reveal substantial differences between locomotor plants of individual fish (***Figure 3***A). Despite these differences, the estimated FRFs shared a common structure: the locomotor dynamics of each fish acted as a low-pass filter on the movements of the nodal point. This common structure enables the application of parametric models, which reduce the complexity of analysis while facilitating computational manipulations of the model system. We adopted the physics-based parametric model of locomotor dynamics of *Eigenmannia* described by ***Sefati et al.*** (***2013***):

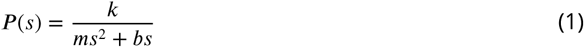

**Figure 3.**
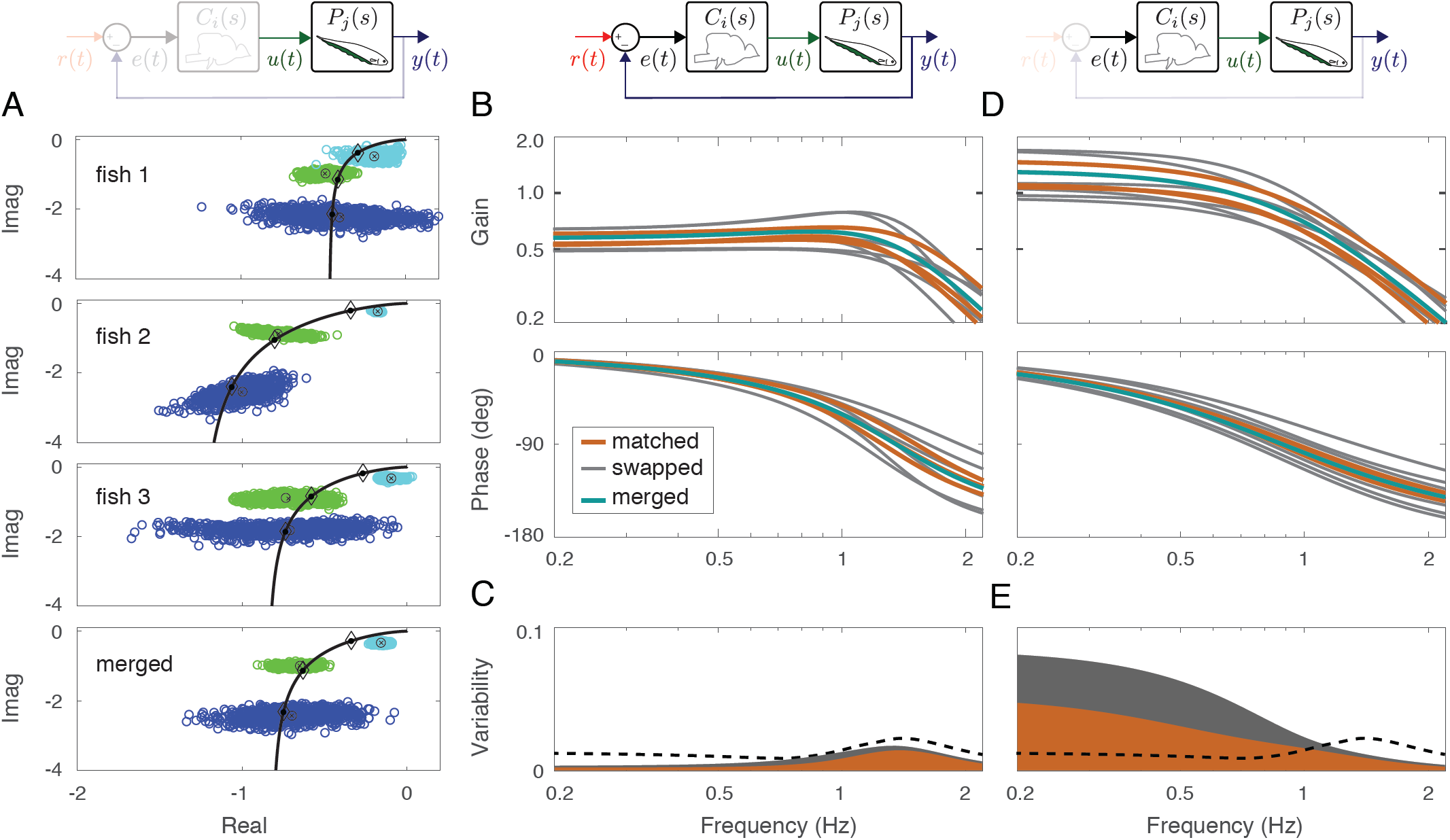
Effects of feedback on behavioral variability. At the top of each column is a control diagram that corresponds to all 1gures in that column. (**A**) Nyquist plots of the estimated plant models for each individual fish and the merged fish. Black lines represent the response of the estimated model with diamonds indicating their values at the test frequencies. Blue, green and cyan point clouds correspond to bootstrapped data estimates for the low (0.55 Hz), medium (0.95 Hz) and high (2.05 Hz) frequencies, respectively. (**B**) Bode gain and phase plots of the estimated input–output models for the matched (orange), swapped (gray), and merged (green) fish in a feedback control topology. (**C**) Variability of the matched (orange) and swapped (gray) models. Dashed line is the trial-to-trial variability observed in the tracking data across fish. (**D**) Same as in (B) but for the loop gain, *P* (*s*)*C*(*s*). (**E**) Same as in (C) but for the loop gain, *P* (*s*)*C*(*s*).

Here, *m, k*, and *b* represent mass, gain, and damping, respectively, and *s* is complex frequency (see, e.g. ***Roth et al.*** (***2014***)). We estimated the parameters in the parametric plant model for each fish based on their FRFs (***Figure 3***A, black lines) via numerical optimization (see Materials and Methods) (***Figure 3–Figure Supplement 1***). Finally, we estimated the plant for a ‘merged’ fish, in which the data from the three fish were concatenated. The differences in the FRFs between individuals resulted in substantial differences (about twofold) in estimated model parameters (***Table 1***). Moreover, the merged fish has plant dynamics that differ from each of the individual fish (***Figure 3***A, bottom), highlighting the need to use individualized plants for the analysis of the control system of each fish. Despite the differences in plant dynamics, fish produced remarkably similar tracking performance, consistent with previously published reports (***Cowan and Fortune, 2007***; ***Roth et al., 2011***).

**Table 1.** Estimated parameters of the plant model for each fish as well as the merged fish.

|  | *fish 1* | *fish 2* | *fish 3* | *merged* |
| --- | --- | --- | --- | --- |
| $k$ (N/m) | 0.3228 | 0.2200 | 0.1622 | 0.2385 |
| $b$ (Ns/m) | 0.0417 | 0.0186 | 0.0218 | 0.0269 |
| $m$ (kg) | 0.0025 | 0.0025 | 0.0025 | 0.0025 |

### Examining the Effects of Feedback on Behavioral Variability

Behavioral robustness could be achieved via precise tuning between the controller and plant dynamics of each fish. Alternatively, the central nervous system (CNS) may be implementing robust feedback control without the need for precise tuning. To test these hypotheses, we built feedback control models that permit computational manipulation of the relationships between controller and plant. We used the second order model proposed by ***Cowan and Fortune*** (***2007***) to represent the input–output behavioral response of the fish:

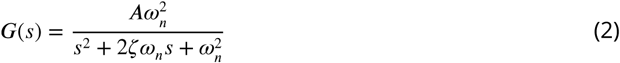

We estimated the parameters for each fish using their movement response *y*(*t*) to refuge movements *r*(*t*). In other words, we generated individualized parametric transfer functions that capture the input–output behavioral performance of each fish. Parameters varied by about 15–20% (***Table 2***).

We investigated how the variability in plant dynamics (twofold) is mitigated at the level of behavior (15-20%). Specifically, we back-calculated a controller for each fish using models of their respective plant dynamics and input–output behavioral responses. Given the plant *P*(*s*) and behavioral performance *G*(*s*) of each individual fish, back-calculation yields a feedback controller for each fish in the following form (***Cowan and Fortune, 2007***):

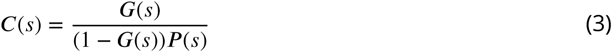

**Table 2.** Estimated parameters of the second order input-output model for each fish as well as the merged fish.

|  | *fish 1* | *fish 2* | *fish 3* | *merged* |
| --- | --- | --- | --- | --- |
| $A$ | 0.53 | 0.60 | 0.52 | 0.57 |
| $\xi$ | 0.58 | 0.55 | 0.52 | 0.55 |
| $\omega_n$ (rad/s) | 8.55 | 9.51 | 7.88 | 8.50 |

Estimates of controllers and plants for individual fish allowed us to implement computational manipulations in the system models. To evaluate the fitness of a given computational model for explaining the biological data, we defined a metric termed ‘trial-to-trial variability’, which provides a conservative estimate of the behavioral variability observed across trials in the real fish. See Materials and Methods (Trial-to-trial Variability) for details. Trial-to-trial variability across frequencies can be seen in ***Figure 3***C,E (black dashed lines).

Having defined the ‘trial-to-trial variability’, we first computed the tracking performance of each fish using its own controller and plant (‘matched’, ***Figure 3***B). Unsurprisingly, the variability of the matched models was lower than the trial-to-trial variability (***Figure 3***C, orange region versus dashed line). See Materials and Methods (Model Variabilty) for details.

To test the hypothesis that the animals rely on precise tuning between their plants and controllers, we substituted the controller of each fish with the plant dynamics of other fish (***Figure 3***B), i.e. a simulated brain transplant (‘swapped’). If the controllers and plants need to be co-tuned, then we would expect a significant increase in variability in the swapped models. Surprisingly, however, the variability remained below the trial-to-trial variability (***Figure 3***C, gray region versus dashed line). These results highlight the fact that sensory feedback can attenuate the output variability despite mismatch between the controller and plant pairs. In other words, feedback models do not require precise tuning between the controller and plant to achieve the low variability we observed in the behavioral performance of the animals.

Having established that output variability is robust to the relations between the plant and controller in a feedback system, we examined the role of feedback. This was achieved by examining the loop gain, i.e. the dynamic amplification of signals that occurs in feedback systems (***Figure 3***D). For our model, this was calculated as the product of the plant and controller in both matched and swapped cases. This removal of feedback revealed dramatic variability in the loop gain at frequencies below about 1 Hz (***Figure 3***E). This variability was well above the trial-to-trial variability observed in fish. In contrast, at frequencies above 1 Hz, the variability was slightly reduced. These results suggest that sensory feedback attenuates behavioral variability in the biologically relevant range of tracking frequencies at a cost of slightly increased variability at high frequencies.

### Parametrizing the Range of Neural Controllers

As predicted by ***Cowan and Fortune*** (***2007***), each of the feedback controllers obtained above for the averaged fish responses had high-pass filtering characteristics despite the differences in their dynamics (***Figure 4***A). What is the range of neural controllers that, when used in this feedback control topology, leads to behavior that is indistinguishable from the real fish? In other words, how well tuned to the plant does the neural controller need to be to achieve these behavioral performances?

**Figure 4.**
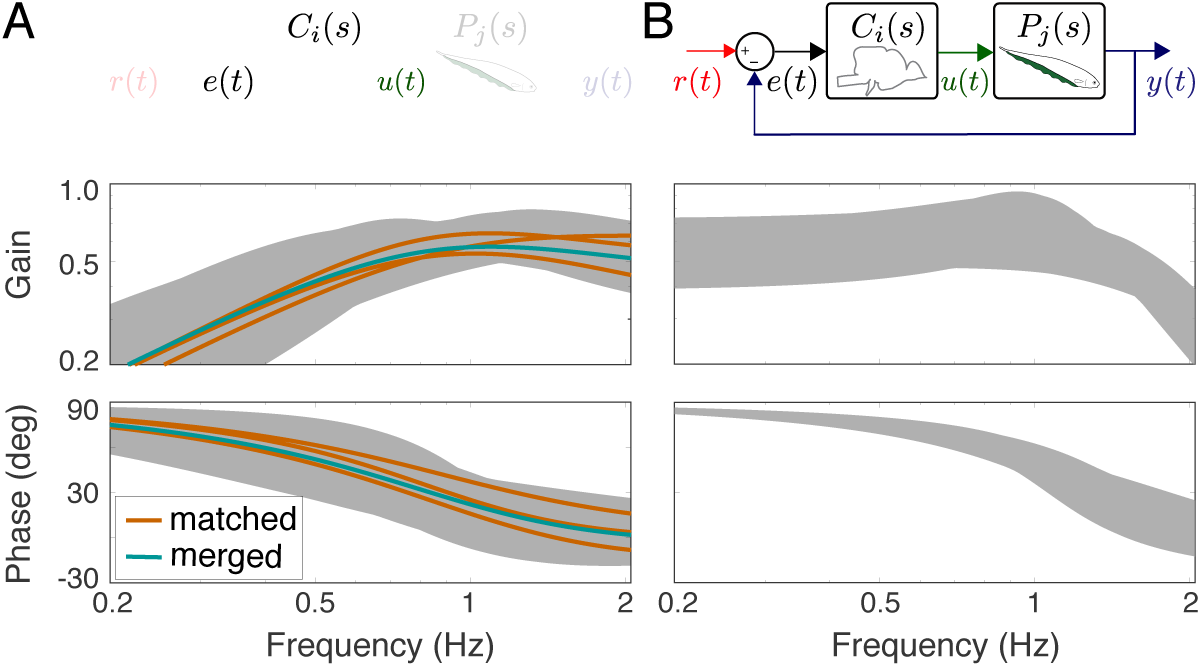
Feedback controllers satisyfing the trial-to-trial variability. (**A**) Bode plots for the estimated controllers for the matched (orange) and merged (green) fish under feedback control. Each controller exhibits high-pass filter behavior across the frequency range of interest. Gray shaded regions represent the range of controllers that produce behavioral responses consistent with trial-to-trial variability. (**B**) The range of behavioral input–output transfer functions consistent with trial-to-trial variability.

We used the Youla-Kučera parametrization to obtain a range of controllers that generate similar behavioral responses (***Kučera, 2011***). Specifically, this parametrization provided a parametric transfer function describing all stabilizing controllers for the merged plant:

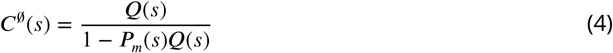

Here, ***P***_*m*_(*s*) is the transfer function of the merged plant and ***Q***(*s*) is any stable and proper function.

***Equation 4*** parametrizes all stabilizing controllers for the merged plant ***P***_*m*_(*s*). However, we were interested in finding the subset of controllers that yields indistinguishable behavioral performances from the real fish. To achieve this, we computed the range for the input–output system dynamics, ***G***(*s*), of real fish response. Specifically, we calculated the bounds for the gain and phase responses of the 1000 input–output transfer function models estimated while computing the trial-to-trial variability (see Materials and Methods). The gray shaded areas in ***Figure 4***B serve as the range of FRFs that are consistent with trial-to-trial variability.

For each of these 1000 transfer functions ***G***(*s*) that are consistent with the trial-to-trial variability of the fish, we selected ***Q***(*s*) = ***G***(*s*)/***P***_*m*_(*s*) to generate 1000 corresponding controllers using ***Equation 4***. The bounds for gain and phase responses of these 1000 controllers (the gray shaded regions) show the breadth of controllers that, when implemented within the feedback control topology, produce behavioral outputs consistent with the performance of the real fish (see ***Figure 4***A). Note that the controllers calculated in ***Equation 3*** also satisfy the structure of ***Equation 4*** when ***Q***(*s*) = ***G***(*s*)/***P***(*s*) for the associated plant dynamics ***P***(*s*) of each fish.

These results suggest that the neural controllers need not be precisely tuned to their associated plant dynamics. We found a wide range of controllers that the fish could implement to generate consistent behavioral performance, although we note that each of these controllers had high-pass filtering characteristics. A similar robustness analysis can also be performed to obtain a range for locomotor dynamics (such as defining bounds for a parametric uncertanity), which are consistent with the trial-to-trial behavioral variability for a given controller.

## Discussion

Feedback-based task control allows animals to cope with dramatic but nevertheless common variations of their plant dynamics between individuals. Further, individuals can experience variations in plant dynamics over time such as instantaneous increases in weight during feeding (***Van Handel, 1965***; ***Hou et al., 2015***), muscle fatigue during repetitive behaviors (***Enoka and Stuart, 1992***), or carrying heavy objects (***Zollikofer, 1994***). These changes likely result in variable mismatch between the neural controller and the locomotor plant of individual animals. This mismatch is similar to that induced by swapping plants and controllers across individuals, suggesting that moment-to-moment variability can also be eliminated through sensory feedback.

Deciphering the interplay between the task plant, behavioral performance, and neurophysiological activity requires understanding the impacts of the closed-loop control topology. Given the range of morphophysiological features observed across individuals within a species, our results suggest that there is also a range of controller dynamics—ultimately manifest as neurophysiological activity—that each individual could use to achieve consistent biologically relevant behavioral performances. As a consequence, we expect to see more variation at the neurophysiological level than is revealed by task performance for behaviors that rely on closed-loop control.

### Reconciling Data-Driven and Physics-Based Models of Locomotor Dynamics

A key contribution of this work is the identification of a data-driven plant model for the locomotor dynamics of a freely behaving animal based on behavioral observations only. To achieve this, we adopted a grey-box system identification approach that seeks to reconcile a physics-based parametric transfer function model with a non-parametric data-driven model (i.e., the frequency-response function).

Developing a model from first principles, e.g. Newton’s laws, is sometimes an effective modeling approach for describing the dynamics of a physical system. For instance, a widely used model in legged locomotion is the spring-loaded inverted pendulum (SLIP) model for describing running dynamics in the sagittal plane (***Blickhan and Full, 1993***; ***Full and Koditschek, 1999***). While physics-based models have proven to be successful in modeling the dynamics of biological and mechanical movements, there are limitations. Physics-based approaches for modeling behaviors at lower levels (e.g., the spiking activity of all motor neurons) may lead to a very complex model that does not accurately capture high-level behavior.

Data-driven system identification approaches are used to directly identify a dynamical model based on empirical data (***Ljung, 1998***; ***Kiemel et al., 2011***; ***Uyanik et al., 2019a***). In general, data-driven system identification may take a black-box approach in which only a general model structure is assumed (say, an ODE or frequency response function). However, data-driven techniques typically generate numerical transfer function estimates to represent animal behavior.

Alternatively, the so-called grey-box approach that we adopt in this paper, integrates the structure of a specific physics-based model—but leaves its parameters free—with data-driven system identification—to fit those parameters. In this case, prior knowledge about the underlying dynamical model informs and constrains data-driven system identification. Grey-box identification can provide a bridge between top-down, data-driven modeling and bottom-up, physics-based modeling. We utilized the parametric dynamical model of ***Sefati et al.*** (***2013***) for the plant but estimated the model parameters using data-driven system identification techniques. Our results show that the data-driven estimates for the plant dynamics match the structure of this model (***Figure 3***A).

### Effects of Variability in Plant Dynamics

Our results reveal two complementary perspectives on variability in plant dynamics. On the one hand, estimates of the closed-loop controllers were highly sensitive to the dynamics of the plant of individual fish. This was an inevitable consequence of our strategy for estimating the controllers—back-calculating the controllers from the plant and closed-loop dynamics—rather than, in and of itself, refuting the precise tuning hypothesis. On the other hand, our closed-loop control models were robust to variability of either the plant or controller, indicating that precise tuning is not needed for this behavior. A control-theoretic sensitivity analysis demonstrates that these results are not unique to this example but rather are a general property of feedback control systems (see ***Csete and Doyle (2002)*** for a review).

Specifically, consider the frequency dependent sensitivity function of the feedback controller *C*(*s*) with respect to plant *P*(*s*), in the closed-loop topology:

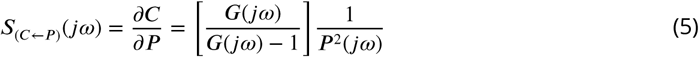

The sensitivity of the controller to the plant dynamics is a frequency dependent function. It depends on the gain and phase of both the measured closed-loop transfer function ***G***(*jω*) as well as the plant model ***P***(*jω*). At low frequencies, fish track extremely well and thus ***G***(*jω*) − 1 ≈ 0. At high frequencies, the low-pass plant ***P***(*jω*) is small. Combining these factors, we expect the sensitivity |***S***(*jω*)| to be large across frequencies. In other words, there is an inescapable sensitivity to plant dynamics when the controllers are estimated using this computational strategy.

We conducted a complementary analysis to compute the sensitivity of the closed-loop tracking response *G* to perturbations in the combined controller and plant dynamics ***PC***. We treated the controller–plant pair ***PC*** as a single variable and obtained the frequency-dependent sensitivity function as

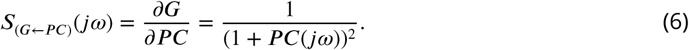

At low frequencies, ***G***(*jω*) has nearly unity gain and thus ***PC***(*jω*) goes to ∞. As a result, sensitivity ***S***_(*G*←*PC*)_ approaches zero. At high frequencies, ***PC***(*jω*) goes to zero and thus sensitivity is bounded around one. Thus, the closed-loop transfer function (in the presence of sensory feedback) is robust against variability of controller–plant pairs despite the fact that the controller estimates are sensitive to plant variations.

### Feedback and Variability in Neural Systems

These findings suggest that a fish could implement a range of controllers in its nervous system for refuge tracking. These controllers must have high-pass filtering characteristics, but their details may be inconsequential. This has two implications for neurophysiological analysis of neural control systems. First, neurophysiological activity within control circuits in open-loop experiments (e.g. playback and anesthetized/immobilized animals) need not appear to be well tuned for the control of a behavioral performance. This poor tuning, which may manifest in variability that appears across levels of functional organization—from variability in neural activity within neurons, variability in tuning across neurons, and variability across individuals—is refined via feedback during behaviors in which the feedback loops are intact. Second, there must be mechanisms by which the controllers are slowly (at time constants greater than that necessary for the moment-to-moment control of the behavior) tuned to the dynamics of the animal’s locomotor plant. For instance, adaptation of cerebellar activity in relation to mismatch between intended versus actual motor performances contribute to the retuning of neural controllers (***Morton and Bastian, 2006***; ***Bell et al., 1997***; ***Pisotta and Molinari, 2014***).

Feedback is mediated both through the effects of behavior on sensory receptors and via descending pathways in the brain. Behavior generally results in concomitant activation of receptor types across the animal, which can include, for example, simultaneous stimulation of stretch receptors embedded in muscles and visual receptors in eyes. Correlations in feedback-related activity across sensory modalities likely contribute to robust control (***Roth et al., 2016***). Internal feedback pathways, interestingly, have been recently shown to synthesize sensory filtering properties of behaviorally-relevant stimuli. Descending neural feedback is used to dynamically synthesize responses to movement (***Metzen et al., 2018***; ***Huang et al., 2018***; ***Clarke and Maler, 2017***). The dynamic synthesis of filters based on current sensory information maybe a mechanism that shapes neural controllers on two time scales: for the moment-to-moment task dynamics and over the longer term for the maintenance of behavioral performances.

## Materials and Methods

All experimental procedures with fish were reviewed and approved by the Johns Hopkins University and Rutgers University Animal Care and Use Committees and followed guidelines for the ethical use of animals in research established by the *National Research Council* and the *Society for Neuroscience*. Adult *Eigenmannia virescens*, a species of weakly electric Gymnotiform fish, were obtained through commercial vendors and housed in laboratory tanks. Tanks were maintained at a temperature of approximately 27° C and a conductivity between 50 − 200 *µ*S. We transferred individual fish to the experimental tank about 1 day before experiments for acclimation. Three fish were used in this study.

### Experimental Apparatus

The experimental apparatus is similar to that used in previous studies (***Stamper et al., 2012***; ***Biswas et al., 2018***; ***Uyanik et al., 2019b***). A refuge machined from a PVC pipe with a length of 15 cm and 5.08 cm diameter was placed in the experimental tank with the fish. The bottom face of the refuge was removed to allow video recording from below. Six windows, 0.625 cm in width and spaced within 2.5 cm intervals, were machined onto each side to provide visual and electrosensory cues. During experiments, we actuated the refuge using a linear stepper motor with 0.94 *µ*m resolution (IntelLiDrives, Inc. Philadelphia, PA, USA) driven via a Stepper motor controller (Copley Controls, Canton, MA, USA). MATLAB (MathWorks, Natick, MA, USA) scripts were used to control the movement of the refuge and to capture video. Video data were captured using a pco.1200s high speed camera (Cooke Corp., Romulus, MI, USA) with a Micro-Nikkor 60 mm f/2.8D lens (Nikon Inc., Melville, NY, USA). All videos used for data analysis were shot at 30 frames per second with 1280 × 1024 pixel resolution. Some videos of ribbon fin motion were shot at 100 frames per second.

### Experimental Procedure

Refuge movement consisted of single sine waves of amplitude 0.1 cm and of frequencies 0.55, 0.95, and 2.05 Hz. The amplitude of refuge movements was chosen because fish rely on counter propagating waves for tracking in this regime (***Roth et al., 2011***). At higher amplitudes, fish often will use a uni-directional wave in the ribbon fin for locomotion. The frequencies were selected to be within the normal tracking regime as determined in previous studies (***Stamper et al., 2012***; ***Biswas et al., 2018***; ***Uyanik et al., 2019b***). Trials were randomized with respect to frequency. Each trial lasted 60 seconds. The stimulus amplitude was linearly ramped up over the first ten seconds to prevent startle responses from the fish. During the experimental phase, the stimulus frequency and amplitude were maintained for 40 seconds. Finally, the stimulus amplitude was ramped down during the final ten seconds. Trials were separated by a minimum break of 2 minutes.

### Reconciling Data-Driven and Physics-Based Approaches to Estimate the Locomotor Dynamics

The position of the refuge and fish were tracked for each video using custom software implemented in MATLAB. The videos were analyzed to extract 3 to 10 seconds segments, where the fish used counter propagating waves for refuge tracking. Then, the nodal point was hand clicked in these video segments: 18000 nodal point measurements were made over a total of 106 segments of data. The physics-based plant model in ***Sefati et al.*** (***2013***) was previously validated with quasi-static open-loop experiments. Here we reconciled the physics-based plant model from ***Sefati et al.*** (***2013***) (***Equation 1***) with the data that were collected in tracking experiments.

For each frequency of refuge movement, M segments of nodal point data were extracted. Each segment of data consists of the following measurements: nodal point shift 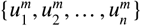 and fish position 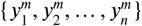 as a function of time 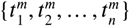, where n is the number of samples and *m* = {1, 2, …, ***M***}.

We estimated the magnitude and phase of the plant model for each frequency of refuge movement. The average value of nodal point shift and fish position were computed from M data segments per fish for each frequency of refuge movement. We aligned each data segment based on the phase of refuge signals. The segments are not completely overlapping: we selected the largest time window with at least 50 percent overlap of data segments. A sine wave function of the following form was fitted to the average nodal point data, *u*_*avg*_(*t*), and average fish position data, *y*_*avg*_(*t*), as

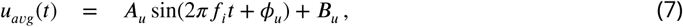

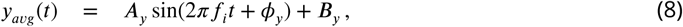

where input–output pairs (***A***_*u*_, ***A***_*y*_), (*ϕ*_*u*_, *ϕ*_*y*_) and (***B***_*u*_, ***B***_*y*_) correspond to magnitudes, phases and DC offsets in polar coordinates, respectively. Note that this fitting was done separately for each refuge frequency, *f*_*i*_ = {0.55, 0.95, 2.05} Hz.

After computing the magnitude and phase for both the average nodal shift and fish position, we estimated the magnitude and phase for the plant transfer function at *ω*_*i*_ = 2*πf*_*i*_ as

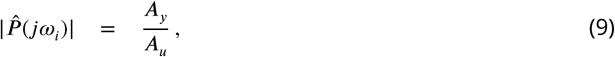

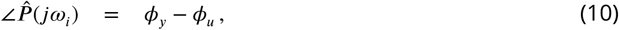

We obtained a non-parametric estimate of the plant transfer function for each frequency *ω*_*i*_, i.e. 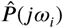 by estimating magnitude and phases. We used 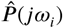 to estimate the parameters of the transfer function model given in ***Equation 1***. In this model, there are three unknown parameters, namely *m, k*, and *b*. However, for the fitting purposes we reduced the number of unknown parameters to two by normalizing the “gain” (*k*) and “damping” (*b*) by dividing by “mass” (*m*). The normalized plant transfer function in Fourier domain takes the form

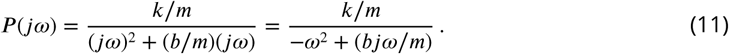

For an ideal deterministic system, for each frequency 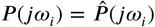, where 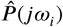 corresponds to the non-parametrically computed frequency response function. For this reason, estimates of transfer function parameters were made by minimizing a cost function using gradient descent method:

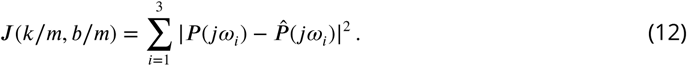

### Trial-to-trial Variability

We examined variability on a trial-to-trial basis for the three individual fish at the three test frequncies. Specifically, across all three fish, we made 37 observations of the frequency response at *f*_*i*_ = 0.55 Hz, 35 observations at *f*_*i*_ = 0.95 Hz, and 34 at *f*_*i*_ = 2.05 Hz, namely:

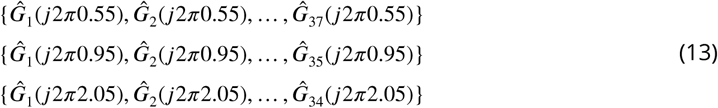

To estimate the trial-to-trial variability at frequencies that were not explicitly tested, we used a parametric approach. Specifically, we constructed 1000 triplets by randomly selecting one FRF from each of the test frequencies in ***Equation 13***. For each of the 1000 triplets, we estimated a transfer function of the form in ***Equation 2*** by using Matlab’s transfer function estimation method ‘tfest’. The variability of the behavioral responses were computed from 0.2 Hz to 2.05 Hz by evaluating the fitted parametric transfer function at each frequency and computing the largest singular value of the central covariance matrix as described in Variability Metric. In addition, the range of the gain and phase of these 1000 transfer function models was plotted in ***Figure 4***B.

### Model Variabilty

The variability across “matched” and “swapped” models was calculated using the frequency responses for both closed-loop transfer function, ***G***(*s*), (***Figure 3***B) and loop gain, ***P*** (*s*)***C***(*s*)), (***Figure 3***D). For the “matched” case, we calculated the variability across the three “matched” models. Similarly, for the “swapped” case, we calculatd the variability across the six “swapped” models. Specifically, variability of these models were computed from 0.2 Hz to 2.05 Hz by evaluating the frequency response functions of the associated matched and swapped models (described below). ***Figure 3***C,E illustrate the variability of the “matched” and “swapped” models both for closed-loop control and loop gain.

### Variability Metric

The variability metric was used to compute trial-to-trial and model variability. The largest singular value of the central covariance matrix of a set of frequencies responses at a given frequency was used as a metric for variability. Specifically, let ***G***_*i*_(*jω*_0_) be a distinct (complex-valued) frequency response function of the fish’s refuge tracking response at *ω*_0_ for a trial *i*, ∀*i* ∈ {1, 2, …, *N*}. Variability across these ***N*** trials at *ω*_0_ was calculated as follows. Let *x*_*i*_ and *y*_*i*_ be real and imaginary parts of the complex-valued frequency response function, namely *G*_*i*_(*jω*_0_) = *x*_*i*_ + *jy*_*i*_. The covariance matrix for the estimated frequency response function in the complex domain was calculated as

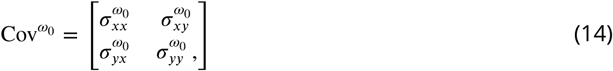

where

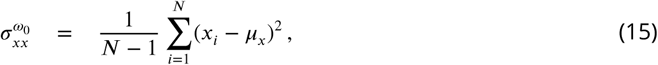

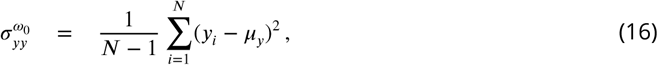

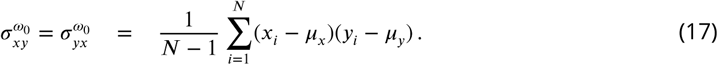

Here, ***µ***_*x*_ and ***µ***_*y*_ are mean values of *x*_*i*_ and *y*_*i*_, ∀*i* ∈ {1, 2, …, *N*}, respectively. Variability was calculated as the largest singular value of the central covariance matrix, 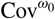.

## Acknowledgments

This work was supported by a collaborative National Science Foundation (NSF) Award to Noah J Cowan (1557895) and Eric S Fortune (1557858). Figures and Figure Supplements incorporate illustrations drawn by Eatai Roth. We also thank him for his invaluable feedback on the manuscript.

---

**Figure 2–Figure supplement 1.**
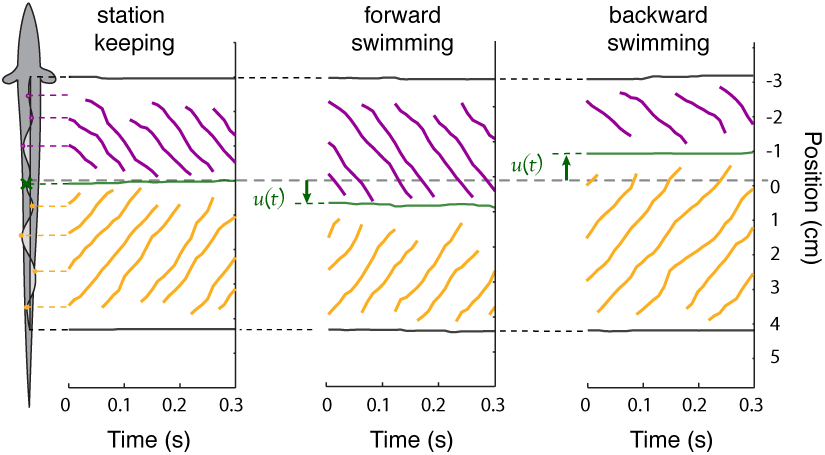
Steady-state position of the nodal point. Three examples of steady-state, constant-speed swimming that illustrate how the peaks and troughs of the ribbon fin wave propagate toward the nodal point. The purple lines indicate the positions of the peaks and troughs of the traveling waves initiated at the rostral end of the ribbon in. Orange lines indicate the traveling waves initiated at the caudal end of the ribbon in. These waves meet at the nodal point (green). (left) Peaks, troughs, and nodal point reference location during station keeping, *dy*/*dt* = 0. (center) The nodal point shifts caudally relative to the reference position during steady-state forward swimming, *dy*/*dt >* 0 and (right) shifts rostrally for backward swimming, *dy*/*dt <* 0.

**Figure 2–Figure supplement 2.**
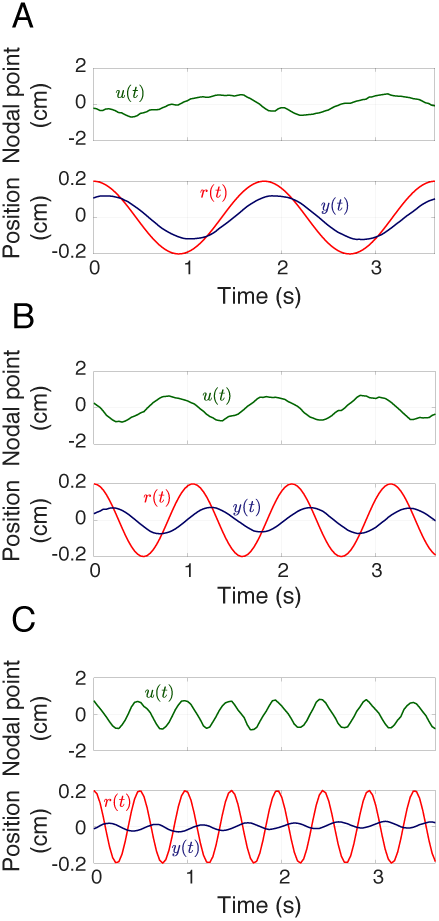
Movement of the refuge, fish and nodal point. Averaged movements (across all trials for one fish) of the nodal point *u*(*t*) (green) and the fish *y*(*t*) (blue) with respect to refuge movements *r*(*t*) (red) at **A**) 0.55 Hz, **B**) 0.95 Hz, and **C**) 2.05 Hz.

**Figure 3–Figure supplement 1.**
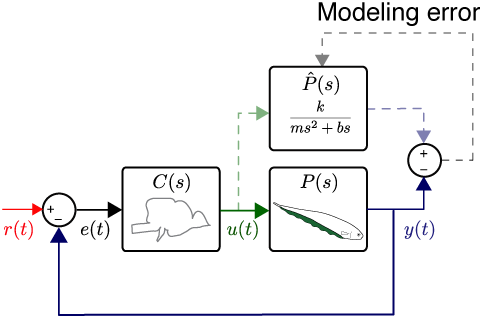
Reconciliation of physics-based and data-driven models. The solid lines represent the natural feedback control system used by the fish for refuge tracking. The dashed lines represent copies of signals used for parametric system identification. 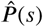 represents the parametric transfer function for the plant dynamics of the fish with “unknown” system parameters. The parametric system identification estimates these parameters via minimizing the difference (modeling error) between the actual output of the fish *y*(*t*) and the prediction of the model.

